# Diversity and specificity of molecular functions in cyanobacterial symbionts

**DOI:** 10.1101/2024.03.22.586251

**Authors:** Ellen S. Cameron, Santiago Sanchez, Nick Goldman, Mark L. Blaxter, Robert D. Finn

## Abstract

Cyanobacteria are globally occurring photosynthetic bacteria notable for their contribution to primary production and their production of toxins which have detrimental impacts on ecosystems. Beyond this, cyanobacteria can form mutualistic symbiotic relationships with a diverse set of eukaryotes, ranging from land plants to fungi. Nevertheless, not all cyanobacteria are found in symbiotic associations suggesting symbiotic cyanobacteria have evolved specializations that facilitate host-interactions. Photosynthetic capabilities, nitrogen fixation, and the production of complex biochemicals are key functions provided by host-associated cyanobacterial symbionts. To explore if additional specializations are associated with such lifestyles in cyanobacteria, we have conducted comparative phylogenomics of molecular functions and of biosynthetic gene clusters (BGCs) in 977 cyanobacterial genomes. Cyanobacteria with host-associated and symbiotic lifestyles were concentrated in the family Nostocaceae, where eight monophyletic clades correspond to specific host taxa. In agreement with previous studies, symbionts are likely to provide fixed nitrogen to their eukaryotic partners. Additionally, our analyses identified chitin metabolising pathways in cyanobacteria associated with specific host groups, while obligate symbionts had fewer BGCs. The conservation of molecular functions and BGCs between closely related symbiotic and free-living cyanobacteria suggests that there is the potential for additional cyanobacteria to form symbiotic relationships than is currently known.

## 1. Introduction

Cyanobacteria, a group of photosynthetic bacteria, have a long evolutionary history with fossil evidence dating back up to 1.9 billion years ago^1^. These organisms are found globally in diverse habitats, from aquatic to terrestrial landscapes and from polar to tropical climates^1–3^. Cyanobacteria have evolved adaptations for survival under numerous types of stresses including desiccation, extreme temperatures, salinity, UV radiation and pathogenic infections^4^. Additionally, they can threaten ecosystems and human health through their production of potent toxins when blooms contaminate water sources^5^. As such, much of cyanobacterial research has focused on public health impacts of freshwater and marine strains^6^. However, the impact of cyanobacteria on ecosystem health extends beyond degrading water quality, as many of the members in this taxonomic group have been found to be critical partners in mutualistic symbiotic associations with a diverse range of eukaryotic hosts.

The keystone example of cyanobacterial symbioses is that of the endosymbiotic event that occurred some 2.1 billion years ago and led to the development of chloroplasts and photosynthetic eukaryotes^7^. Beyond the endosymbiont origin of chloroplasts, cyanobacteria are also found in symbiotic associations with diverse hosts such as protists, metazoans, fungi, macroalgae and land plants in both terrestrial and aquatic environments^7–9^. These symbionts provide hosts with beneficial services including photosynthetic products^10^ and fixed nitrogen^11,12^. The mode of host association is also variable, including epiphytic growth (e.g. on feathermoss), intra-organismal location in specialised symbiotic structures and intracellular incorporation^9,11^. These associations can be ancient with examples of cyanobacterial symbionts found in a fossilised lichen from 400 million years ago ^7^. Such long associations raise the potential for coevolution between the eukaryotic host and specialised cyanobacterial partners^9^, selecting for symbiotic competence^13,14^.

Host-symbiont interactions require pathways for communicating and detecting signals^14^ which may involve secondary metabolites. Secondary metabolites, compounds that are not essential for primary growth and reproduction^15^, are produced by co-localized genes organized as biosynthetic gene clusters (BGC)^16^. These compounds are often specialized for species interaction and survival in stressful environments, and can include bioactive compounds with antibacterial, antifungal and cytotoxic properties^4,15,16^. Secondary metabolites have previously been shown to impact symbiotic associations such as diatoms producing compounds to promote growth and attachment of beneficial bacteria^17^, or coral microbiomes producing a high diversity of antimicrobial products^18^. However, secondary metabolites are often produced for a specific physiological or ecological reasons and are often taxon specific^19^, with this specificity potentially being a mechanism for symbiont communication to their potential host^20^.

Even amongst microbes known for their production of diverse secondary metabolites, cyanobacteria alone are known to produce over 1,100 unique secondary metabolites and their genomes frequently contain a high number of BGCs^21,22^. The majority of cyanobacterial genomes contain polyketide synthase and nonribosomal peptide synthetase pathways that account for up to 5% of their total genome sizes^23^. The compounds that cyanobacteria produce span diverse roles ranging from UV protection (mycosporines and scytonemin) to grazing deterrents and nutrient scavenging^6^ which may provide additional competitive advantages to hosts^24^. The compounds may also mediate inter-partner communication in symbioses. For example, the production of nostopeptolide in the cyanobacterium genus, *Nostoc*, is associated with repression of formation of infectious differentiated cells and is down-regulated in the presence of plant hosts^25,26^. While genome mining approaches have identified many cyanobacterial biosynthetic gene clusters of unknown function^27,28^, the potential for symbiosis-specific secondary metabolites and their distribution among lineages of cyanobacteria has not been fully explored.

Cross-talk between cyanobacteria and their host-species has been previously reported, ranging from the upregulation of transcription of ammonium and nitrate transporters^29^ to influencing cell differentiation in the life cycle of *Nostoc*^11^. However, varying reports of host-specificity and phylogenetic clades of symbiont cyanobacteria^9,11,30–32^ requires a phylum-wide study to explore the origins of host-association in this ancient lineage. Uniquely, *Nostoc* has shown broad symbiotic competence with different eukaryotic hosts, yet there still remains questions on molecular drivers of these associations due to the potential of non-host specific responses as isolates from cycads have previously been shown to also enter into symbiotic associations with mosses, fungi and angiosperms (*Gunnera*)^13^. Previous research has identified niche-specific BGCs that have been connected to individual host-specific associations in cyanobacteria^33^ suggesting the presence of specialized secondary metabolites associated with cyanobacterial symbionts. However, a large-scale analysis of all available cyanobacterial genomes within the context of symbiotic associations has not yet been conducted. In this work, we utilize comparative phylogenomic approaches to identify trends in distribution of (i) molecular functions and (ii) biosynthetic gene clusters which may mediate host-symbiont interactions in this phylum.

## 2. Methods

### 2.1 Cyanobacterial genomes, habitat annotation & quality control

Assembled genome sequence data for 1078 species belonging to the phylum Cyanobacteria were downloaded from NCBI RefSeq in January 2023 (Table S1). An additional 27 metagenomic assembled genomes (MAGs) taxonomically classified as cyanobacteria from lichen sources^34^ were included to provide additional examples of host-associated symbionts for a total of 1105 cyanobacterial genomes.

Wherever possible the sampled cyanobacteria were assigned to their source habitat(s) based on available sample metadata, associated publication(s) or metadata describing the original isolation reported by culture collections. These habitat assignments include aquatic (e.g., freshwater, marine and man-made aquatic sources) and terrestrial (e.g. soils), as well as host-associated environments. Host associations include vascular and non-vascular plants, protists, fungi, macroalgae, and marine mammals (epidermal mats). Individual host species were grouped into broad taxonomic categories including bryophytes, cycads, fruit trees, diatom endosymbionts, and lichens. Water fern (*Azolla*) cyanobacterial symbionts were placed in their own category. These habitat annotations were also used for grouping the cyanobacteria into two broader lifestyle classifications: free-living and host-associated. Cyanobacterial genomes of which no specific source habitat could be discovered were excluded, leaving 1026 cyanobacterial genomes for comparative analyses.

Quality control filtering was performed using CheckM^35^ (Version 1.1.3) and 977 high-quality (>90% complete; <5% contamination; Table S2) cyanobacterial genomes were retained for phylogenetic tree reconstruction and downstream analysis. Representatives of Melainabacteria (n=37), a basal non-photosynthetic lineage of cyanobacteria, were included as an outgroup.

### 2.2 Phylogenetic tree reconstruction

Taxonomic classification of genomes and generation of marker gene alignments was conducted using GTDBtk^36^(v. 2.3.0; Table S3). Phylogenetic trees were constructed for the final high-quality set of cyanobacterial genomes using IQ-TREE^37^(v. 2.2.0). The analysis used the LG+R10 model as identified in the IQ tree model finder based on the Bayesian Information Criterion (BIC). A family-level phylogenetic tree for the family Nostocaceae (n = 300), rooted with representatives of the order Elainellales, was constructed using IQ-TREE and the LG+F+R7 model determined by BIC. Phylogenetic trees were visualised using iTOL^38^(v.5). For phylum and family level trees, non-parametric bootstraps (n = 1000) were conducted with IQ-TREE to assess the robustness of phylogenetic inferences.

### 2.3 Genome annotation and KEGG completeness estimation

Cyanobacterial genomes were annotated with Prokka^39^(v.1.14.6) and the resulting gene predictions were functionally annotated with KofamScan(v.1.3.0) to derive Kyoto Encyclopaedia of Genes and Genomes (KEGG) module annotations. KofamScan predictions were used with KEGG-Decoder^40^(v. 1.3) to generate a table representing molecular function completeness across samples (Table S4). KEGG functions were classified as being present using two thresholds, either >98% complete for a more stringent analysis of distribution and complete function, or >50% complete for lower stringency examination for the potential presence of molecular functions, herein referred to as indicative functions (Figure S1). Presence/absence matrices generated for KEGG functions were used in a phylogenetic logistic regression^41^ to identify enrichment of molecular functions based on lifestyle classification at the phylum level (Table S5, S6) and enrichment of molecular functions in individual isolation sources in the family Nostocaceae (Table S7, S8). Phylogenetic logistic regressions were conducted using the *phyloglm* function in the R package *phylolm*^42^, using the penalised likelihood with Firth’s correction and 100 bootstraps. Responses of lifestyle classification and isolation sources were defined as significant if the p-value was less than 0.05.

### 2.4 Biosynthetic gene cluster prediction and classification

BGCs were predicted on cyanobacterial genomes using SanntiS^43^ (v. 0.9.1) due to high performance on both isolate genomes and MAGs, thus providing consistent annotations across all genome types. The predictions were subsequently filtered to remove those occurring at the edges of contigs and those which were less than 3000 bp in length, reflective of the minimum length of BGCs observed in the MIBiG database^16^. BGCs were initially classified by SanntiS into standard classes such as ribosomally synthesised and post-translationally modified peptides (RiPPs), terpenes, nonribosomal peptides, polyketides, alkaloids, saccharides, and hybrid classes which represent BGCs that cover multiple biochemical classes (Table S9). To detect enrichment of total and specific BGC classes in host-associated symbionts, phylogenetic linear regression was conducted at the phylum level (Table S10) and in the Family Nostocaceae (Table S11). This was performed with the *phylolm* function using 100 bootstraps and a lambda model for covariance.

To expand upon the basic BGC classifications provided by SanntiS and identify diversity in potential products, predicted BGCs in cyanobacteria were clustered with a large, reference set of biosynthetic gene clusters (the ‘reference BGC collection termed RefBGC hereafter). RefBGC includes BGC predictions from running SanntiS on MGnify^44^ and RefSeq genomes^45^, as well as the BGCs found in MiBIG^16^, and subsequently refined to only include complete predictions. This clustreing enabled the assignment of BGCs to more specific groups based on functional domain composition, utilizing the Louvain community detection method^47^ and the Sørensen-Dice similarity coefficient^48^. To refine the SanntiS BGC classification assigned to each group antiSmash^46^(v.7.0.0) predictions were also generated for RefSeq and used to provide more specific natural product annotations, thereby combining the breadth offered by SanntiS and the accurate BGC product assignments provided by antiSMASH. Groups of BGCs containing antiSmash predictions were retained as the final set of BGCs (Table S12). The habitat source of each BGC group was use in phylogenetic logistic regression to identify enrichment of specialized biosynthetic gene clusters in cyanobacteria with different lifestyles (Table S13, Table S14). This was performed with the *phyloglm* function maximizing the penalized likelihood with Firth’s correction across 100 bootstraps. Groups found to be significantly enriched at the phylum level were used to assess phylogenetic signal in the family Nostocaceae using the D-statistic^49^ with the *phylo.d* function in the R package *caper*^50^(v.1.0.2) of lifestyle classification and isolation sources were defined as significant if p-value was less than 0.05.

## 3. Results

### 3.1 Enrichment of Molecular Functions and Biosynthetic Gene Clusters in Host-Associated Cyanobacterial Symbionts

Using the taxonomic classifications based on GTDB the cyanobacterial genomes were assigned to 18 taxonomic orders and 42 families, which were monophyletic based on the GTDBtk phylogeny thus facilitating rigorous interpretation of evolutionary relatedness of these organisms. Of these, Cyanobacteriales (n = 576) and PCC-6307 (representative of *Cyanobium gracile*; n = 261) comprised over 85% of available genome assemblies (Figure 1A). Habitat sources were highly skewed, with aquatic environments (n = 753) representing >75% of environmental sources for all genome assemblies. Notably, only 6% (n = 62) of assessed cyanobacterial genomes were isolated from host-associated environments including non-vascular and vascular plants, protists, macroalgae, metazoan epidermal mats and fungi. Within this, Cyanobacteriales accounted for 93% [5.9% of host-associations in all assessed cyanobacterial genomes; n = 58] of all host-associated cyanobacterial symbiont genomes including representatives from all detected habitat source classifications (Figure 1B). NCBI taxonomy was also considered, however due to challenges with nested, non-monophyletic groupings based on current taxonomic nomenclature, comparisons based on ‘taxonomic identity’ were not possible. Nevertheless, similar trends were shown with NCBI taxonomy with a high proportion of genomes arising from the orders Synechococcales (n = 428) and Nostocales (n = 300) comprising nearly 75% of available reference genome assemblies with host-associations concentrated in the Nostocales (Figure S2).

**Figure 1:**
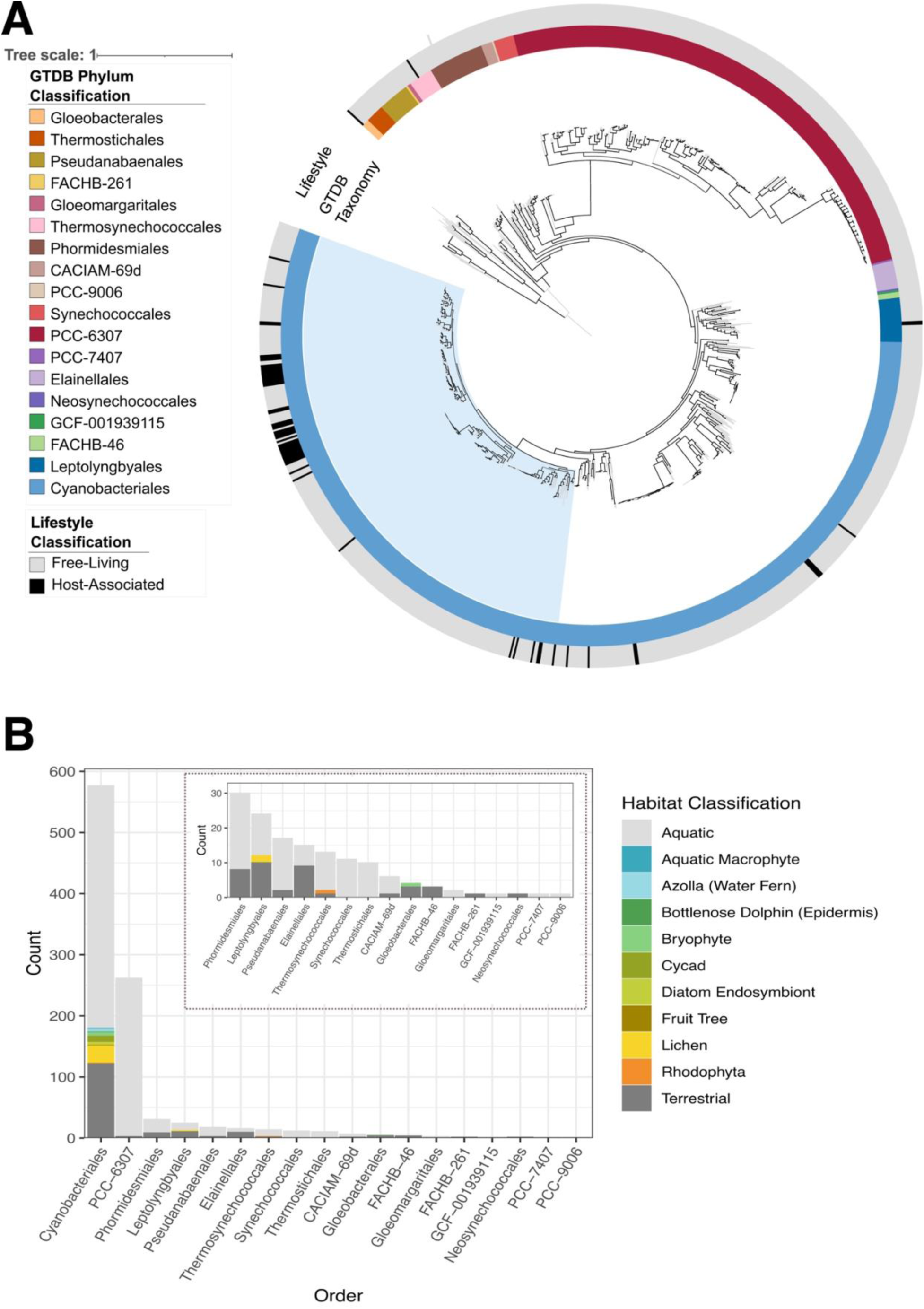
Phylogeny and distribution of host-associated lifestyles in the phylum, Cyanobacteria. (A) Phylogeny generated using concatenated marker genes of genome sequences of strains from phylum Cyanobacteria, rooted with representatives of the sister group, Melainabacteria, with 1000 bootstraps. Branches with high bootstrap support (>80%) are shown with black. The outer annotation track depicts the lifestyle classification to highlight host-associated cyanobacterial symbionts. The inner annotation track depicts the classified taxonomic order assigned by GTDB. Nostocaceae, a family containing the majority of host associations, are shaded in light blue. (B) Frequency counts distributed across taxonomic orders for habitat classifications highlighting the different host sources including vascular plants (water fern, cycad, fruit trees, aquatic macrophytes), non-vascular plants (bryophytes), protists (e.g. diatoms), macroalgae (Rhodophyta), fungi, and epidermal mats of aquatic mammals such as dolphins.

KEGG functional annotations were analysed to identify molecular functions enriched in symbiont genomes by exploring the distribution of complete KEGG functions. In total, 77 complete KEGG functions were variably present across the phylum, of which 20 were significantly associated with lifestyle classification (Figure 2A; Figure S3). Host-associated lifestyles were found to have a significantly higher level of occurrence of functions including those of glucogenesis (p=0.042; Est. 0.85), Fe-Mn transporter (p=5.53e-07; Est. 1.45), glucoamylase (p=4.86e-04; Est. 1.31), zeaxanthin diglucoside production (p=6.36e-04; Est. 1,01), cobalt-magnesium transporters (p=3.46e-02; Est. 0.54) and nitrogen fixation (p=4.77e-02; Est. 0.59). While statistically non-significant, chemotaxis (p=0.056; Est. 0.41) was also found to have a higher likelihood of occurrence in host-associated cyanobacteria. Certain complete functions were also found to have a significantly lower occurence in host-associated symbionts including photosystem II (p=7.97e-14; Est. −2.74), the MEP-DOXP pathway (p=1.87e-11; Est. −2.37), methionine (p=1.24e-09; Est. 3.07) and leucine metabolism (p=2.36e-03; Est. −0.83), F-type ATPase (p=5.54e-06; Est. −2.03), NAD(P) H-quinone oxidoreductase (p=1.25e-05; Est. −1.18), riboflavin biosynthesis (p=3.25e-05; Est. −1.59), sulfide oxidation (p=3.58e-04; Est. −1.17), urea transporters (p=1.68e-03; Est. −0.81), cytochrome bd complex (p=2.87e-03; Est. 0.87), cobalt transport proteins (CbiMQ; p=5.42e-03; Est. 0.79), D-galacturonate epimerase (p=6.21e-03; Est. 0.71), formate dehydrogenase (p=5.36e-03; Est. −1.11), cytochrome bd complex (p=2.87e-03; Est. −0.8375) and iron transport system binding proteins (p=2.56e-03; Est. −0.85). Assessment of indicative functions also revealed a significantly higher likelihood of occurrences for Type I Secretion systems (p=0.024; Est. 0.53) and SecSRP secretion pathways (p=0.01; Est. 0.82).

**Figure 2:**
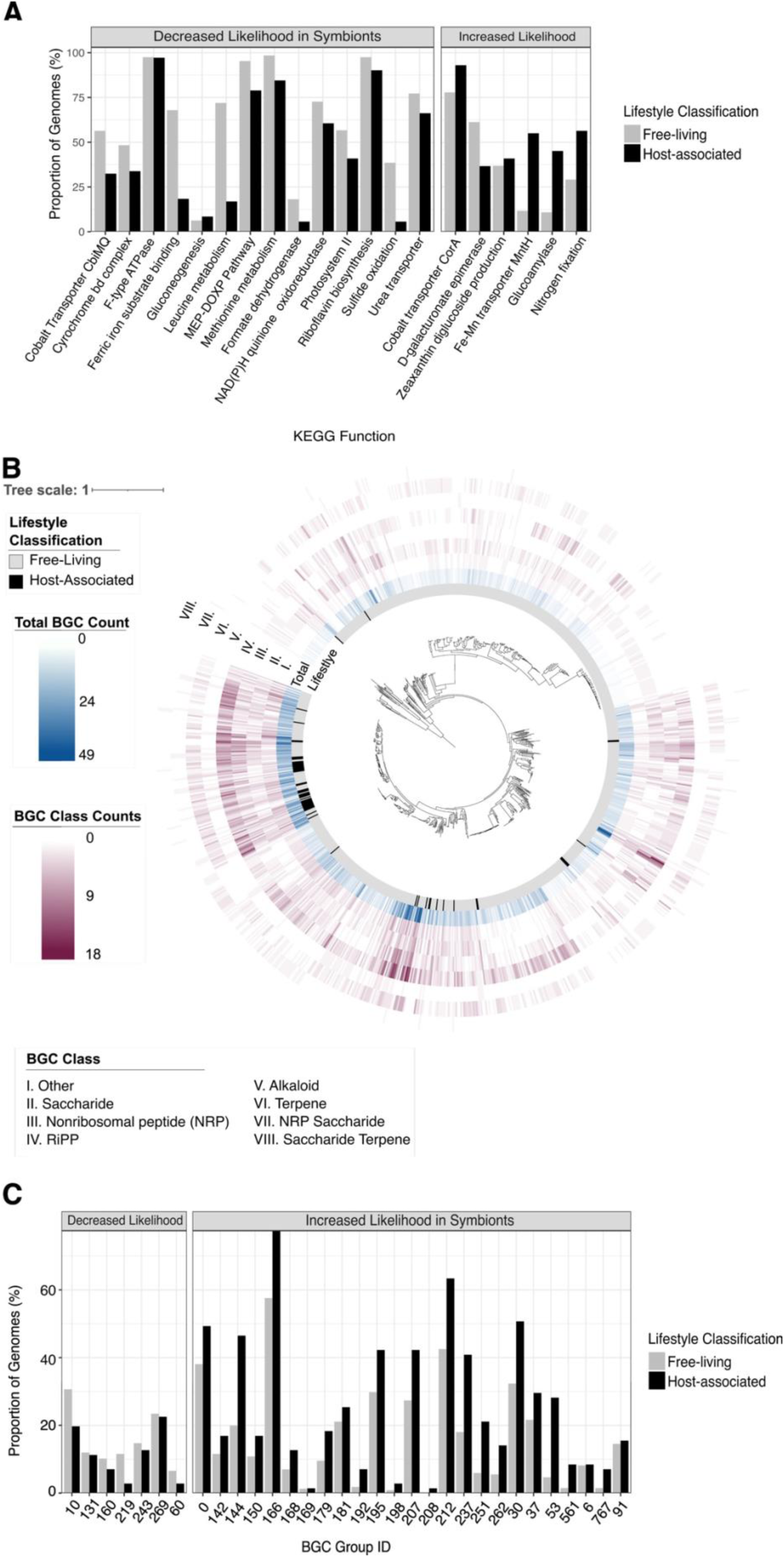
Host-associated enrichment of KEGG pathways and secondary metabolite production potential. (A) Proportion of genomes of each lifestyle type containing KEGG pathways shown to be significantly impacted by life-style classification including key functions corresponding to beneficial ecosystem services including nitrogen fixation. (B) Distribution of counts of total detected biosynthetic gene clusters and classes of biosynthetic gene clusters shown to be significantly impacted by lifestyle classification across the phylum Cyanobacteria (C) Proportion of genomes for each lifestyle type with unique groups of biosynthetic gene clusters that are significantly impacted by lifestyle classification.

To further explore specializations associated with host-associated relationships in cyanobacteria, 8,815 BGCs were identified across 98% (n=961) of all cyanobacterial genome assemblies. In total, 21 classes of biosynthetic gene clusters were identified including hybrid classes which span biochemical properties of multiple classifications. Lifestyle classification was found to significantly associate with the number of detected BGCs. Host-associated cyanobacteria were found to have a significantly lower numbers of BGCs in total (p = 1.02e-06; Est. −4.13) (Figure 2B), and this trend was paralleled at the level of individual BGC class. Host-associated cyanobacterial symbionts were found to have significantly lower count of individual biosynthetic gene cluster classes including nonribosomal peptides (NRP; p = 0.016; Est. −0.52), RiPPs (p = 1.44e-04; Est. −1.21), alkaloids (p=1.01e-03; Est. −0.15), terpenes (p=2.84e-03; Est. −0.43), saccharides (p=8.7te-03; Est. −0.47), saccharide terpenes (p=0.019; Est.-0.038), NRP saccharides (p =0.042; Est. −0.082), and other (p = 0.0012.87e-05; Est. −0.75639), a class of BGC that does not fit into properties of otherwise described secondary metabolites.

The 8815 biosynthetic gene clusters identified were classified into 124 unique groups representative of BGCs, which are likely to produce similar secondary metabolites based on similarity of the protein domain annotations. Although host-associated symbionts were found to have a significantly lower count of BGCs and classes of BGCs as a whole, individual BGC groups were found to be positively associated with cyanobacterial symbionts. Overall, 61 groups were found to be present in both free-living and host-associated cyanobacteria, 61 groups were found only in free-living cyanobacteria, and only 2 groups were found exclusively in host-associated symbionts corresponding to a terpene in a cycad symbiont and ‘other’ classification in aquatic macrophytes symbionts. Of the 61 groups found in both free-living and host-associated cyanobacteria, 25 were found to have a significantly higher prevalence in host-associated cyanobacteria (Figure 2C; Figure S4), while only 7 showed a significantly decreased prevalence in host-associated symbionts (Table S10).

### 3.2 Host-Associated Lifestyle Appears Non-Specific with Multiple Origins in the Nostocaceae

Cyanobacteriales-classified cyanobacteria were recovered as a well-supported monophyly (Figure 1A) and contained the majority of the symbionts analysed. Within the Cyanobacteriales, the host-associated lifestyle was found to be concentrated in the family Nostocaceae (Figure 3A). Phylogenetic reconstruction based on marker genes from publicly available high-quality cyanobacterial genomes belonging to the family Nostocaceae revealed a family-wide distribution of host-associated growth forms (Figure 3B). Eight monophyletic clades corresponding to a unique host category (Table S15) ranging in levels of host specificity. Denoted clades I–VIII, they derive from: diatom endosymbionts; Peltigeraceae lichens *Solorina crocea* and *Peltigera malacea*; the lichen *Peltigera membranacea*; *Azolla* ferns; an unspecified lichen thallus cyanobiont culture ATCC 53789); the lichen *Peltigera*; Peltigeraceae lichens *Collema furfuraceum*, *Leptogium austroamericanum*, *Lobaria pulmonaria*, *Peltigera membranacea*, *Peltigera aphthosa* and *Peltigera malacea*; and *Dioon* cycads, respectively.

**Figure 3:**
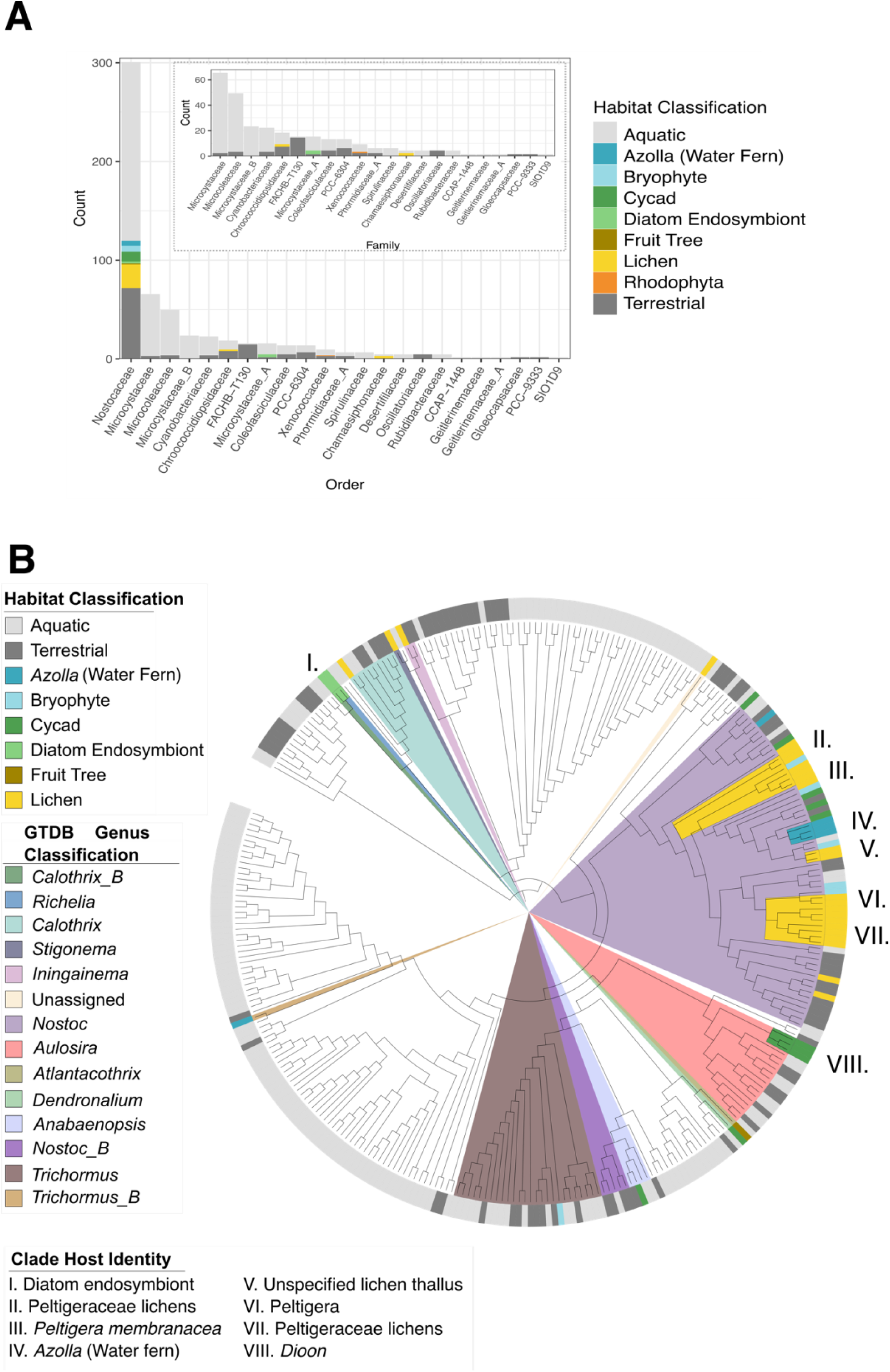
Distribution of host-types in the order Nostocales and the origin of host associations in Nostocaceae. (A) Frequency counts distributed across taxonomic families in the order Nostocales which includes the majority of host-associated cyanobacterial symbiont genomes spanning a high diversity of eukaryotic hosts in the family, Nostocaceae. Families with low frequency counts are displayed as an inset panel. (B) Cladogram of Nostocaceae generated from an alignment of marker genes rooted with the outgroup of Elainellales (n = 15) to explore the origin of host-specific association. Genera with host-associations are highlighted, as well as a non-host associated genus of *Nostoc* (*Nostoc_*B).Colour block shading on branches represent eight monophyletic clades containing symbionts arising from single host classifications.

Ten cyanobacterial genomes were sourced from cycad symbioses but only three of these were found to form a monophyletic clade. *Aulosira*, previously classified as *Nostoc*, comprised monophyletic clade VIII. These symbionts were all from a *Dioon* host supporting previous reports of monophyletic origin of endophytic cyanobacteria with this host species (Guiterrez-Garcia et al., 2019). Cyanobacteria from other cycad hosts (*Cycas revoluta* (n = 3), *Macrozamia* (n = 1), *Zamia pseudoparasitica* (n = 1), *Encphalartos horridus* (n = 1), and *Euterpe edulis* (n = 1)) were distributed across the phylogeny. The genomes sourced from *Cycas revoluta* did not form a monophyletic clade and were distributed across the Nostocaceae tree. The cyanobacterium from the Arecales palm, *Euterpe edulis,* was found in a clade with the cyanobacterium from *Garcina macrophylla,* a dicot (Malpighiales) fruit tree.

Clade IV contained 3 of the 5 analyzed *Azolla* cyanobionts. Notably, the cyanobiont isolated from an epiphytic growth form on *Azolla* was not found with other true *Azolla* cyanobionts.

Five of the monophyletic clades, denoted II, III, IV, V and VII, contained 66% (n = 16) of the analysed lichen cyanobionts, and their hosts were all Peltigeraceae fungi. Lichen cyanobionts most distant to the main lichen clades arose from lichens of different family lineages including *Coccocarpia palmicola* (Coccocarpiaceae) and *Placynthium petersii* (Placynthiaceae) in more basal origins of the Nostocaceae. While all lichens observed in this analysis were of the order Peltigerales, the mycobiont from these lichens are in a different fungal family compared with those in the other analysed cyanolichens (Peltigeraceae), suggesting the potential for genomic diversity in cyanobionts depending on host identity.

Bryophyte cyanobionts did not form host-specific clades, but instead were often found in clades containing lichen cyanobionts or terrestrial isolates. Bryophyte cyanobionts were limited to three host species: *Blasia pusilla* (n=3), *Phaeoceros* (n=1), and *Leiosporoceros dussi* (n=3). The multiple isolates from *Blasia pusila* and *Leiosporoceros dussi* were distributed across the tree but commonly observed in clades with lichen cyanobionts.

### 3.3 Host-specific molecular specialization in Nostocaceae symbionts

To identify host specialization of cyanobacterial symbionts in the family Nostocaceae, the occurrence of KEGG functions across specific isolation sources was assessed. A total of 69 complete KEGG functions were found across Nostocaceae genomes. 5 of these were found in 99% (n=299) of Nostocaceae genomes including functions of histidine, tyrosine and arginine metabolism, nostoxanthin production and retinal biosynthesis. An additional 30 were found in more than 90% of Nostoacaeae genomes with functions including amino acid metabolism, astaxanthin production, starch and glycogen degradation, riboflavin biosynthesis and sulfolipid biosynthesis, and Type I secretion systems. Exploration of indicative functions identified the ubiquitous distribution of many additional functions present in 90% of Nostocaceae genomes, including nitrogen fixation, Sec-SRP secretion pathways, chemotaxis, and cobalamin and thiamine biosynthesis. Some of these ubiquitous functions had also been observed to be significantly enriched in host-associated genomes at the phylum level. In addition to the ubiquitous distribution of certain molecular functions, specific isolation sources were also found to be associated with the prevalence of certain molecular functions (Figure 4A; Table S7,S8; Figure S5). Sulfur dioxygenase (0.027; Est. −2.52) had a significantly lower prevalence in symbionts isolated from the water fern, *Azolla*. Lichen cyanobionts were shown to have functions that had either significantly increased or decreased likelihood of occurrence. Phosphonate transporters (p=0.012; Est. −1.23), methionine synthesis (p=0.026; Est. −1.82) and cytochrome bd complex (p=0.037; Est=-1.03) were found to have a significantly lower prevalence in lichen cyanobionts. Conversly, lichen cyanobionts had significantly higher likelihood for Fe-Mn transporters (p=2.71e-03; Est. 1.48), glucoamylase (p=0.049; Est. 1.03) and photosystem II (p=3.29e-03; Est. 1.61). Similarly, cycads symbionts were also found to have significantly higher likelihood for complete pathways for glucoamylase (p=2.48e-03; Est. 2.23) and chitinase (p=0.021; Est. 1.252), and nearly significant increased likelihood for Fe-Mn transporters (p=0.057; Est. 1.26).

**Figure 4:**
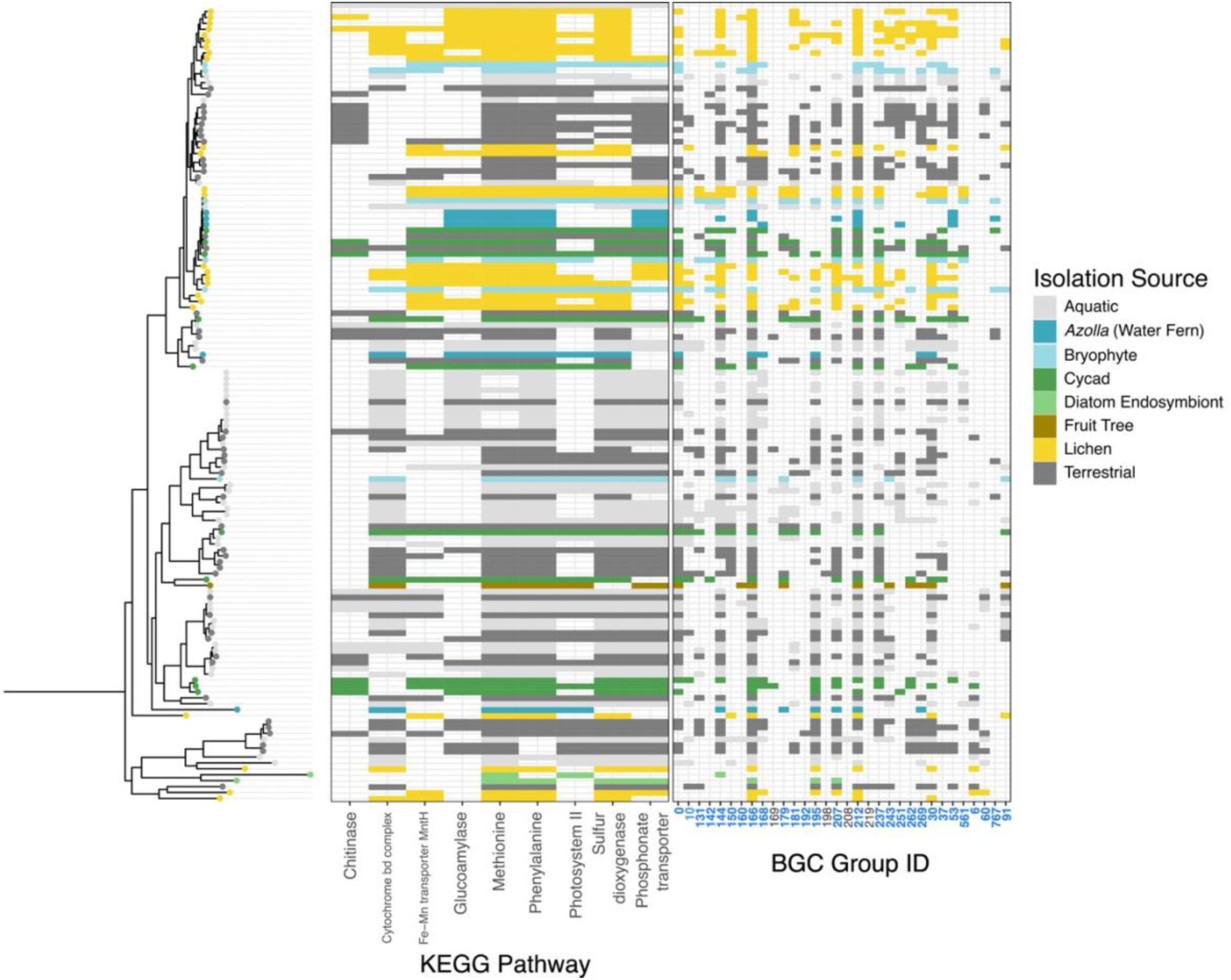
Distribution of significant KEGG functions and groups of biosynthetic gene clusters impacted by isolation source in Nostocaceae genera which include host-associated cyanobacterial symbionts. (A) KEGG functions found to be significantly impacted by specific isolation sources including host-associated symbionts from cycads, lichens and the water fern, *Azolla*. (B) The distribution of 32 BGC groups identified as being significantly impacted by lifestyle-classification (i.e. free-living vs. host-associated in genera of Nostocaceae with host-associate cyanobacterial symbionts. Group names shown in bolded blue font face indicate a significantly non-random phylogenetic distribution indicating shared evolutionary history.

The distribution and occurrence of classes of BGCs in the family Nostocaceae revealed trends correlated with host identity (Table S11; Figure S6). The cyanobacterial symbionts of the water fern, *Azolla*, were found to consistently have a significantly lower number of total BGCs(p=5.19e-05; Est. −12.32), nonribosomal peptides (p=2.54e-03; Est. −2.26), nonribosomal peptide polyketides (p=2.54e-03; Est. −1.53), RiPP (p=8.45e-03; −2.86), terpenes (p=9.72e-03; Est. −1.25), ‘other’ (p=9.54e-04; −2.43), a class of BGC that does not fit into properties of otherwise described secondary metabolites. Other symbionts were also found to have a significantly lower number of BGCs including fruit tree symbionts with a significantly lower number of non-ribosomal peptides (p=0.046; Est. −3.85), saccharide terpenes in cycad symbionts (p=0.03; Est. −0.095) and lichen cyanobionts with a significantly lower number of RiPPs (p=2.84e-03; Est. −2.09). In addition to reduced counts, some symbionts were found to have significantly increased numbers of terpenes (p=0.0.19; Est. 1.00), alkaloid terpenes (p=0.016; Est. 0.16), and NRP polyketides (p=8.64e-03; 1.33) in bryophytes symbionts, and polyketide saccharides (p=4.88e-29; Est. 1.00)in fruit tree fruit tree symbionts.

All 32 groups of BGCs that were shown to be significantly impacted by lifestyle classification were detected in the family Nostocaceae (Figure 4B; Figure S7). Of these, 28 groups had a significantly non-random evolutionary-distribution, and those which had non-significant phylogenetic signal (groups 169, 198, 208 and 219) were sparsely present within this family. 21 BGC groups were identified to be significantly impacted by specific isolation source with a significantly increased prevalence being observed commonly in multiple terrestrial host-associated environments (e.g., cycad, lichen, bryophytes) alongside free-living terrestrial cyanobacteria.

## 4. Discussion

We have compiled and analysed a large dataset of high-quality cyanobacterial genomes to explore the distribution of taxa that are associated with eukaryotic hosts, and to investigate the biochemical diversity and commonalities that distinguish symbionts and free-living isolates. These features could be observed broadly at the phylum level in both molecular functions (as predicted through KEGG orthologs) and BGCs. Broadly, these specialized functions can be summarized into 4 key categories: nitrogen fixation, carbohydrate utilization, environmental communication, and mediation of biotic interactions via secondary metabolite production. We both confirm some of the current understanding of cyanobacterial symbiotic associations and identify novel host specific features in symbiont genomes.

The provision of fixed nitrogen to their eukaryotic hosts is one of the key benefits of cyanobacterial symbiosis in both plant^9,11,12^ and lichen systems^51,52^. We found enrichment of nitrogen fixation in host-associated cyanobacterial symbionts across the phylum and ubiquitously in the family Nostocaceae, supporting this as one of the key mutualistic beneficial services. Nitrogen fixation in cyanobacteria requires iron^53^ and has also been shown to require manganese in legume nodule bacterial symbionts^54,55^, and we demonstrated increased occurrence of Fe-Mn transporters in host-associated cyanobacteria at the phylum level and in cycad and lichen symbionts within the family Nostocaceae.

Carbohydrate-active enzymes including chitinase, glucoamylase and L-lactate dehydrogenase were found to have a significantly higher prevalence in host-associated cyanobacterial symbionts. Notably, in the family Nostocaceae, chitinase was only found to have a significantly higher prevalence in cycad symbionts. Chitin, a highly abundant polysaccharide, is a key component in the cell walls of fungi ^56,57^ and may serve as a source of nitrogen for cyanobacterial and algal growth ^56^. The presences of carbohydrate utilization genes in bacteria are related to the habitats they are isolated from, with enrichment of carbohydrate metabolism correlated with the carbohydrate composition of the environment^58^. The potential for microbes to target the fungal cell wall to prevent pathogenic fungal infection of plant hosts^57^ suggests a potential additional mutualistic benefit of the cyanobacterial symbionts found in cycads. The relative absence of chitinase activity loci in lichen symbionts demonstrates a potential selection against antifungal activity and a key difference in fungal versus plant-cyanobacterial symbioses. While the other enriched carbohydrate-active enzymes observed at the phylum level were not found to be enriched in specific host types, it will be interesting to explore in more detail the trends in distribution of carbohydrate active enzymes in cyanobacteria to align these results with patterns previously reported across the prokaryotic tree of life^58^.

With the exception of diatom endosymbionts and the water fern, *Azolla*^11^, the majority of cyanobacterial symbionts are not permanently associated with the host. Thus, cyanobacterial symbionts require the ability to sense and locate hosts. This may be achieved through chemotaxis involving signal transduction pathways in response to chemical attractants produced by plants^59^ and the ability to sense chemoattractants has proven to be critical in the formation of plant symbioses^59,60^. Consideration of partially complete KEGG functions revealed chemotaxis to have a higher prevalence in host-associated cyanobacteria, but is not significant (p=0.057, near significant). This function was also observed across the Nostocaceae taxa correlating with the occurrence of host-associated symbionts. The enrichment of motility functions has also been previously reported in terrestrial cyanobacteria^61^. As the majority of these symbiotic associations, especially true of those found in terrestrial systems, are facultative for the cyanobacteria^9,11^, this raises the important question of whether free-living cyanobacteria that possess these characteristics are also potential symbiotic partners and whether the diversity of symbiotically competent cyanobacteria is significantly higher than currently reported.

In addition to the ability to sense and respond to their environment, two secretion systems (Type I secretion systems and Sec-SRP) were also found to have a significantly higher likelihood of occurrence in host-associated symbionts suggesting specialization to release products into the environment. While other secretion systems are known to be used to colonize hosts for pathogenic and symbiotic activity (e.g., Type III secretion systems transporting product directly into a eukaryotic cell)^62^, Type I secretion systems are capable of transporting products to the extracellular space in a single step^63^. As observed in bacteria that promote plant growth, the benefit of these microbial partners is often dependent on the secretion systems^64^. However, in the case of the cyanobacterial symbionts, the questions of what beneficial and symbiotically critical compounds may be produced and released by these organisms and how they vary depending on the eukaryotic host remains unexplored.

One of the most notable patterns in the distribution of classes of biosynthetic gene clusters was observed in Nostocaceae symbionts of the water fern, *Azolla*. These symbionts consistently had a significantly lower number of total BGCs, which was paralleled in specific classes including nonribosomal peptides, nonribosomal peptide polyketides, RiPPs,terpenes, and ‘other’. Cyanobacterial symbionts of *Azolla* represent the only currently known permanent obligate symbionts^11^. As secondary metabolites, particularly terpenes, often have roles in mediating complex ecological interactions^6^, so the reduced BGC content in these obligate symbionts may be representative of the reduced complexity of their environment. As *Azolla* symbionts are permanently associated with their host, the requirement for response to environmental stress and to mediate interactions with other organisms is reduced in comparison to cyanobacterial symbionts located in facultative mutualisms where they also need to survive as free-living bacteria.

Reduced numbers of RiPPs were observed in lichen symbionts. RiPPs have very diverse functions ranging from quorum sensing to antifungal and antibacterial properties^65^. Metagenomic sequencing of lichens has forced a reconceptualisation of the symbiosis from a one mycobiont-one photobiont model to one that encompasses additional fungal partners and a diverse microbiome^34,66^. This diversity may play a critical role in the growth of the lichen^34^. That lichen cyanobionts have fewer RiPPs may reflect adaptation to coexistence in this diverse community, and is a topic worthy of deeper analysis.

In contrast to overall reduced counts of biosynthetic gene clusters, symbionts in bryophytes and fruit trees were found to have increased numbers of BGCs predicted to produce terpenes, alkaloids, nonribosomal peptides, and polyketide saccharides. These BGC systems may be responsible for important ecological interactions^18^. Examination of specific unique groups of BGCs in the family Nostocaceae notably revealed that these groups occur in both free-living and host-associated cyanobacteria, and are often not restricted to individual host types. We note that this pattern contrasts previous research suggesting niche specific BGCs only in cycad symbionts^33^. Cyanobacterial isolates from cycads have also been shown to be capable of forming symbiotic associations in laboratory conditions with mosses, mycorrhizal fungi and *Gunnera* (a flowering plant)^13^. This supports our findings of the potential of unspecific host symbiotic competence in secondary metabolite profiles as demonstrated by our large-scale analyses of cyanobacteria and cyanobacterial symbionts.

Previous phylogenetic reconstruction of Cyanobacteria has presented contrasting conclusions concerning the relationships of symbiotic isolates: (i) proposing clades that are comprised of cycad, bryophyte and lichen symbionts^32^; (ii)separation into clades representative of extracellular or intracellular/extracellular symbionts^9^; (iii) grouping of lichen symbionts^67^; or (iv) grouping of plant-associated symbionts^33^. We found host-associated cyanobacteria were scattered across the phylogeny, with few monophyletic clades of symbionts, as previously reported for Nostoc isolates from lichen symbionts^31^. Monophyletic clades of cyanobionts involved in symbioses were detected in isolates from diatom endosymbionts, *Dioon* cycads, sets of Peltigeraceae lichens and the water fern, *Azolla*. In Nostocaceae the basally arising host-associated samples corresponded to lichen symbionts associated with the fungal families Coccocarpiaceae and Placynthiaceae. The other Nostocaceae lichen symbionts analysed were associated with fungal family Peltigeraceae, and were placed intermixed with free-living, *Azolla*-associated and bryophyte-associated isolates. As the lichen fungal partner is known to display a preference in photobiont acquisition^68,69^, it may be that Coccocarpiaceae and Placynthiaceae fungi have a different range of potential partners than the Peltigeraceae. It will be highly informative to generate genomic data for additional, diverse cyanolichens.

In many cyanobacterial symbioses the symbiont may be found in a host association or as a free-living form: these life habits are not mutually exclusive. The availability of free-living cyanobacteria in surrounding environments influences the symbiotic partners found in host associations^11,70^ and free-living cyanobacteria closely related to symbiont clades may prove to be potential symbiotic partners. The increased prevalence of specific BGCs observed across both free-living cyanobacteria in terrestrial environments and symbionts found in terrestrial host-associations (e.g., lichens, cycads, bryophytes) further demonstrates this potential for an increased diversity in cyanobacterial symbionts than has currently been observed. Future research focused on generating novel cyanobacterial genomes from additional symbiotic associations will be critical in advancing the understanding of host range and symbiont diversity in the phylum Cyanobacteria.

## Data Availability

The data analysed during in this study are available from RefSeq and the European Nucleotide Archive (ENA) repositories with accession numbers provided in Supplementary Table S1.

## Acknowledgements

This research was supported by EMBL (European Molecular Biology Laboratory) core funds.

