## Supplementary figures for "Diversity and specificity of molecular functions in cyanobacterial symbionts"

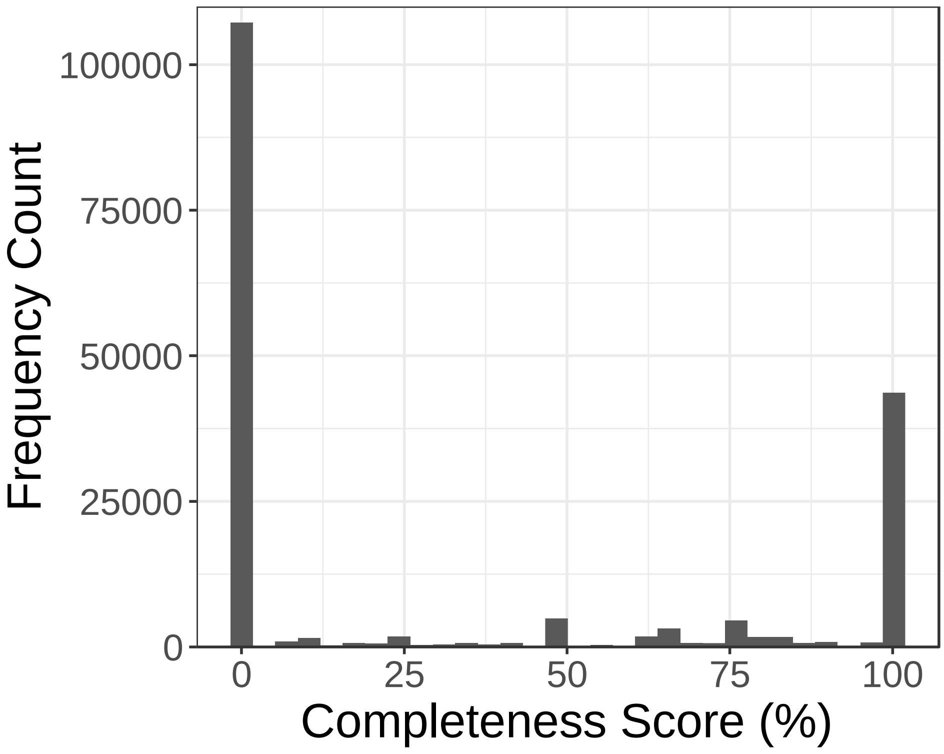


**Figure S1:** Frequency count of KEGG functions completeness score. Majority of detected functions occur in high completeness.

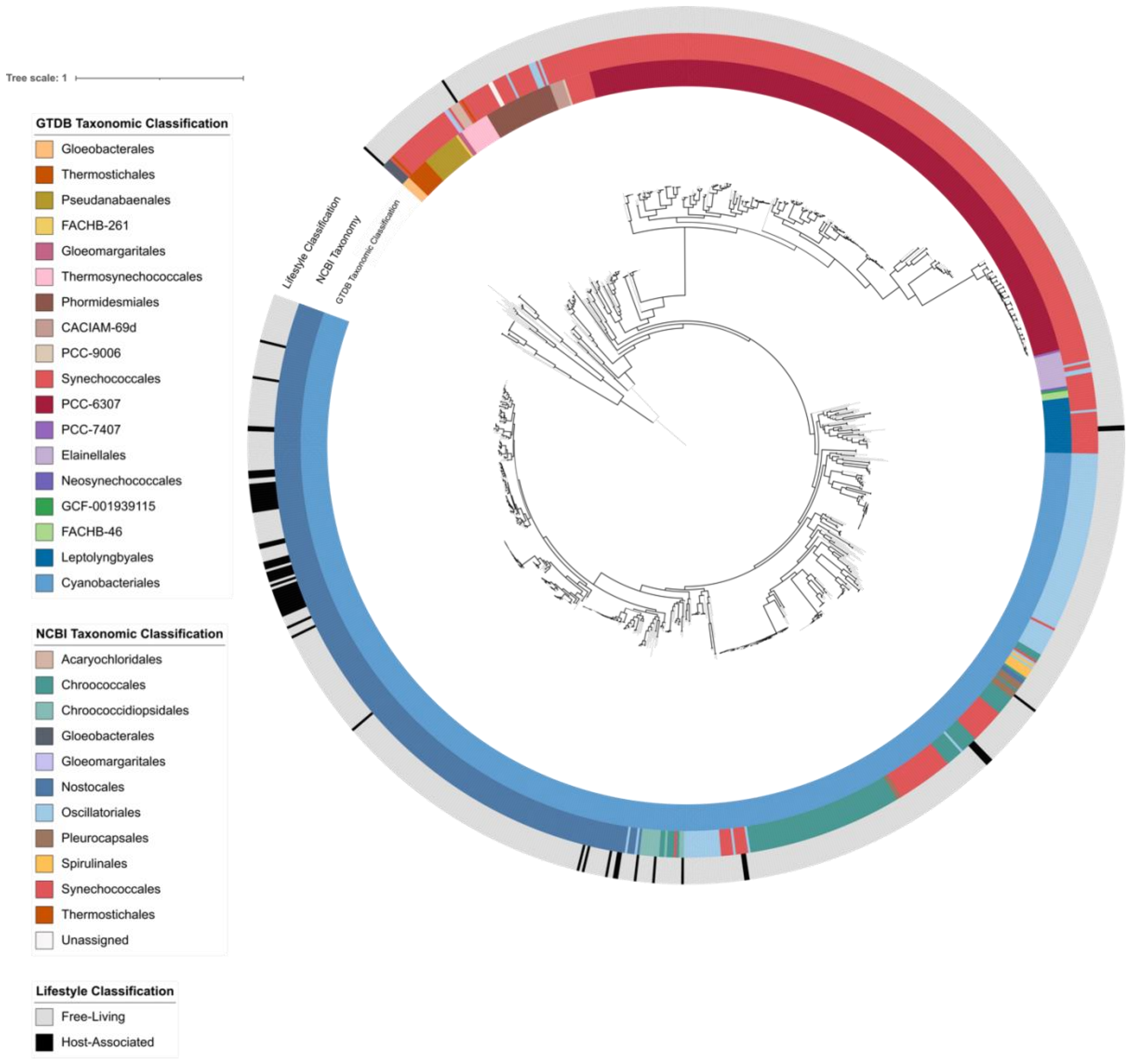


**Figure S2:** Phylogeny generated using concatenated marker genes of genome sequences of strains from phylum Cyanobacteria, rooted with representatives of the sister group, Melainabacteria, with 1000 bootstraps. Branches with high bootstrap support (>80%) are shown with black. The outer annotation track depicts the lifestyle classification to highlight host-associated cyanobacterial symbionts. The inner annotation track depicts the classified taxonomic order assigned by GTDB, and the middle track depicts the NCBI taxonomic classification revealing non-monophyletic distribution of current orders across the phyla.

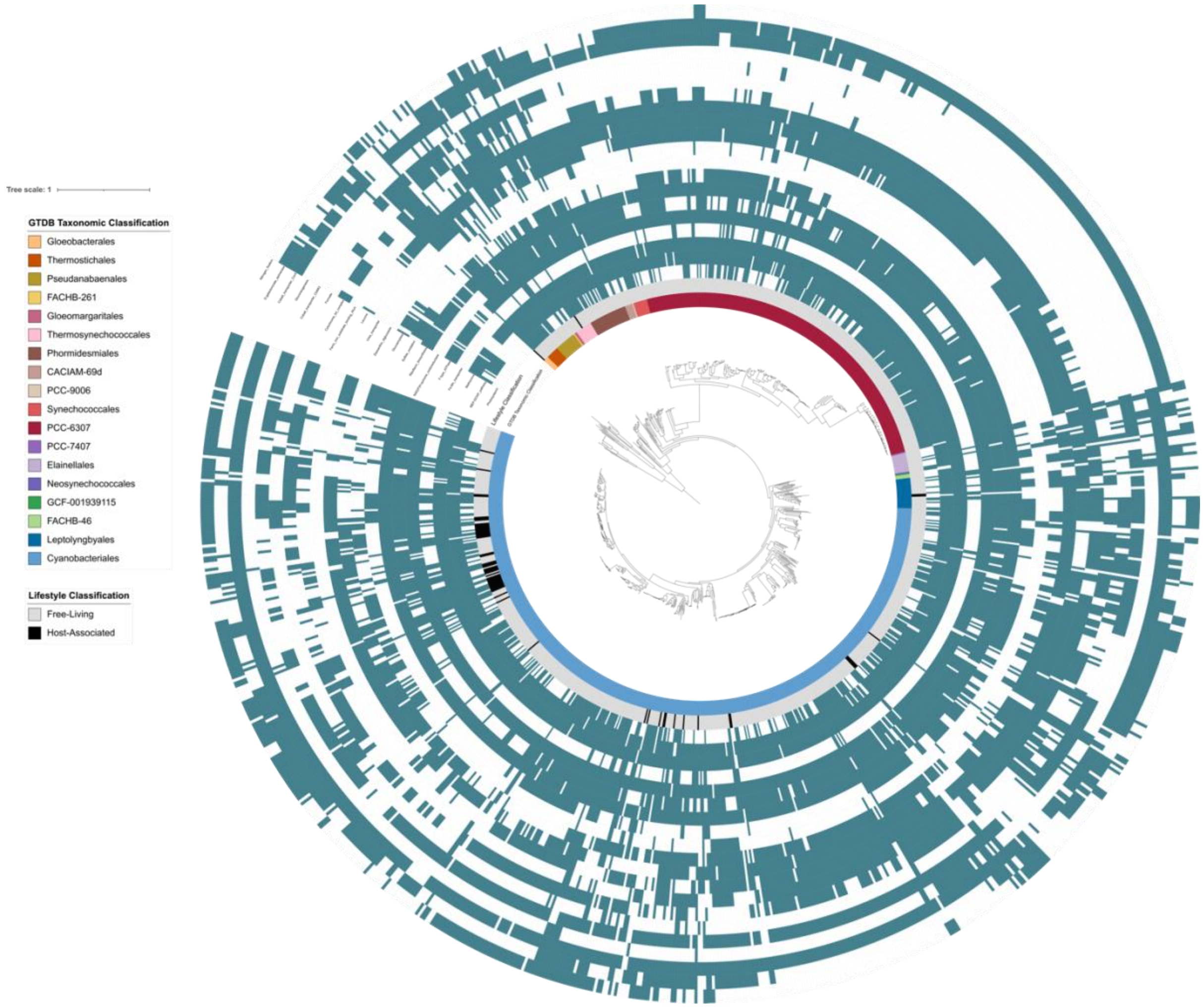


**Figure S3:** Phylogeny generated using concatenated marker genes of genome sequences of strains from phylum Cyanobacteria, rooted with representatives of the sister group, Melainabacteria, with 1000 bootstraps. Branches with high bootstrap support (>80%) are shown with black. From inside to outside annotation tracks depict the i) classified taxonomic order assigned by GTDB, ii) lifestyle classification to highlight host-associated cyanobacterial symbionts, and iii) remaining tracks depict the distribution of molecular functions found to be significantly associated with lifestyle classification.

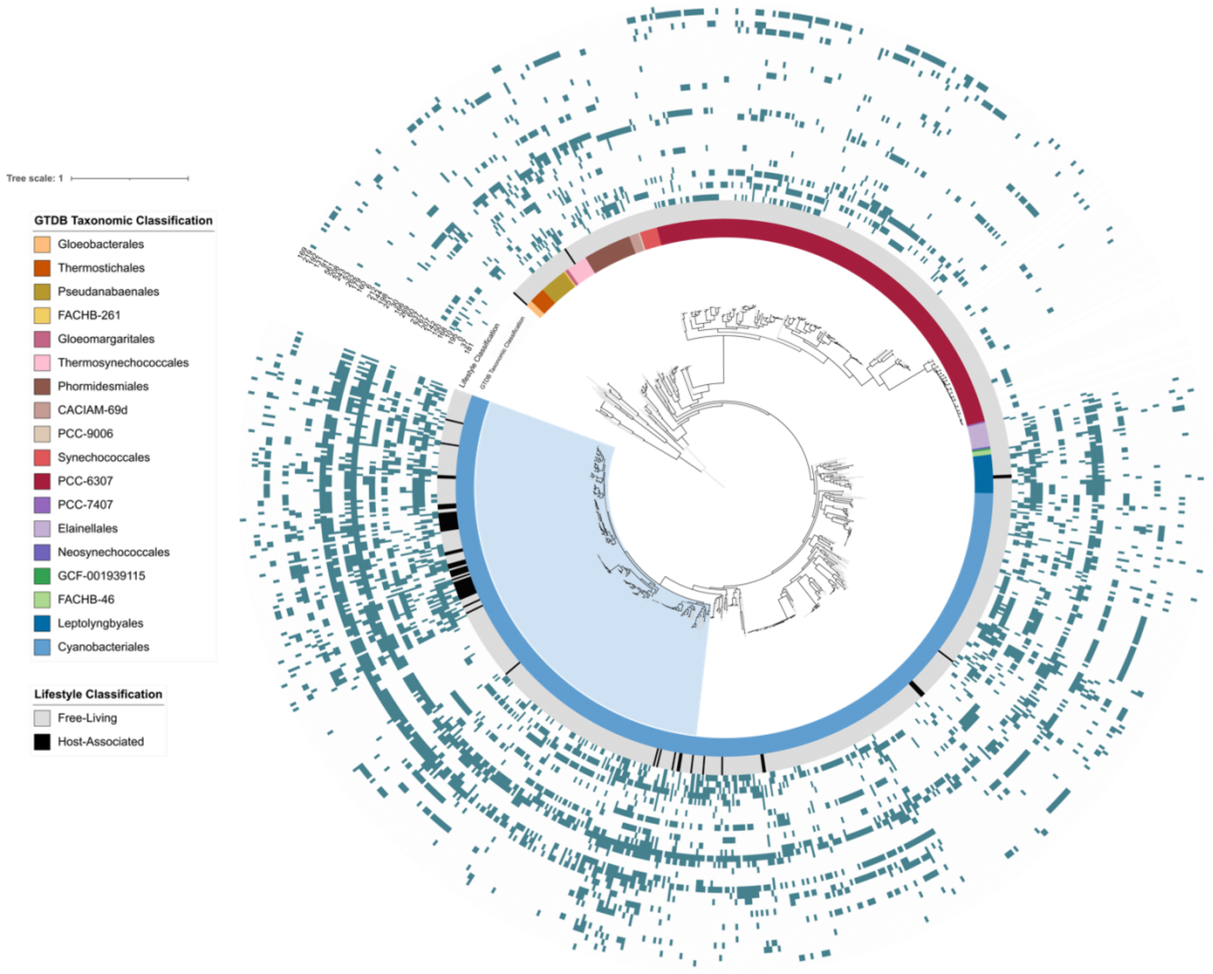


**Figure S4:** Phylogeny generated using concatenated marker genes of genome sequences of strains from phylum Cyanobacteria, rooted with representatives of the sister group, Melainabacteria, with 1000 bootstraps. Branches with high bootstrap support (>80%) are shown with black. From inside to outside annotation tracks depict the i) classified taxonomic order assigned by GTDB, ii) lifestyle classification to highlight host-associated cyanobacterial symbionts, and iii) remaining tracks depict the distribution of BGC groups generated through Louvain4 clustering found to be significantly impacted by lifestyle classification.

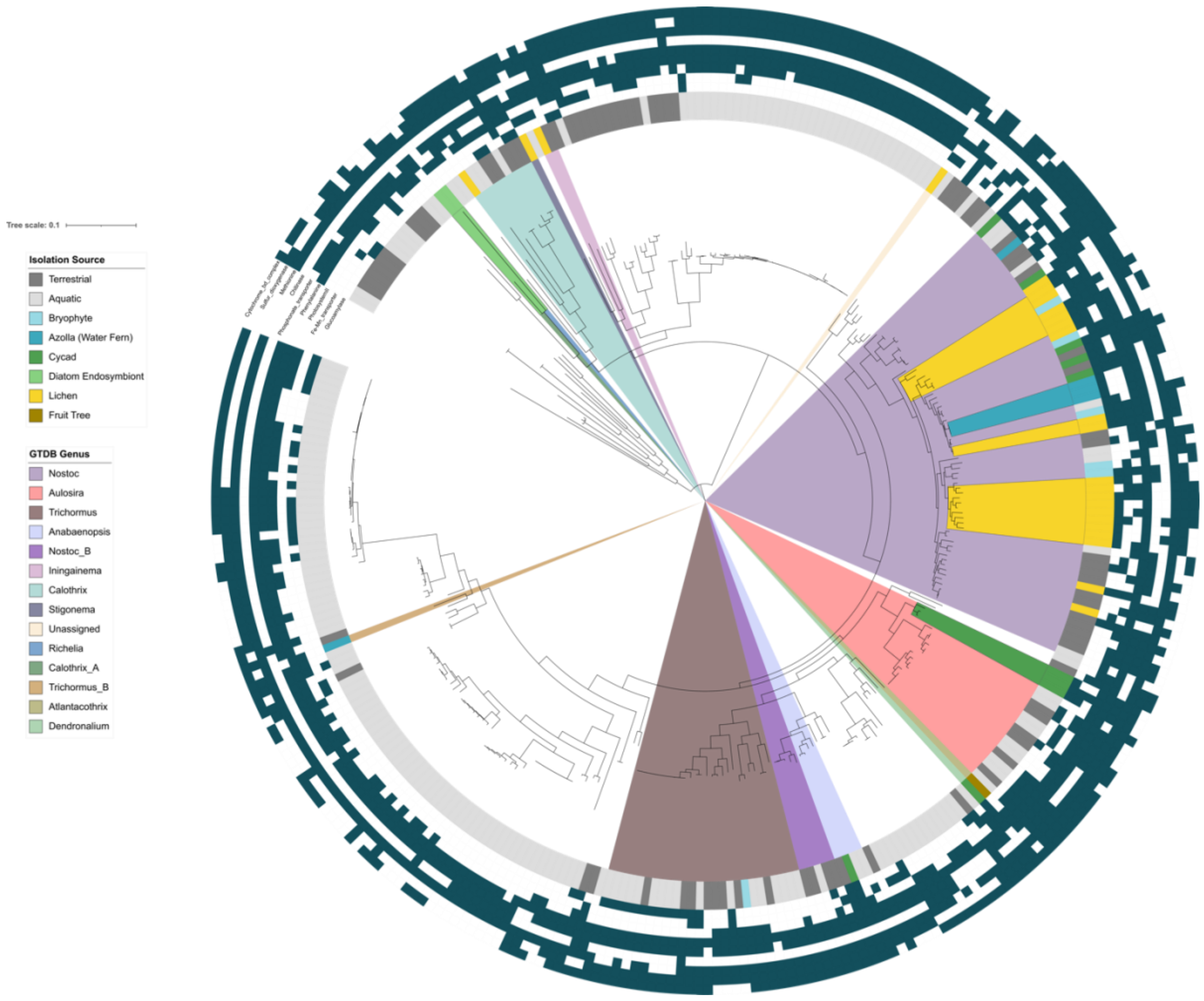


**Figure S5:** Phylogeny generated using concatenated marker genes of genome sequences of strains from family Nostocaceae, rooted with representatives of the order Elainellales, with 1000 bootstraps. Genera with symbiotic lifestyles are highlighted. From inside to outside annotation tracks depict the i) isolation source, and ii) remaining tracks depict the distribution of molecular functions found to be significantly associated with isolation source.

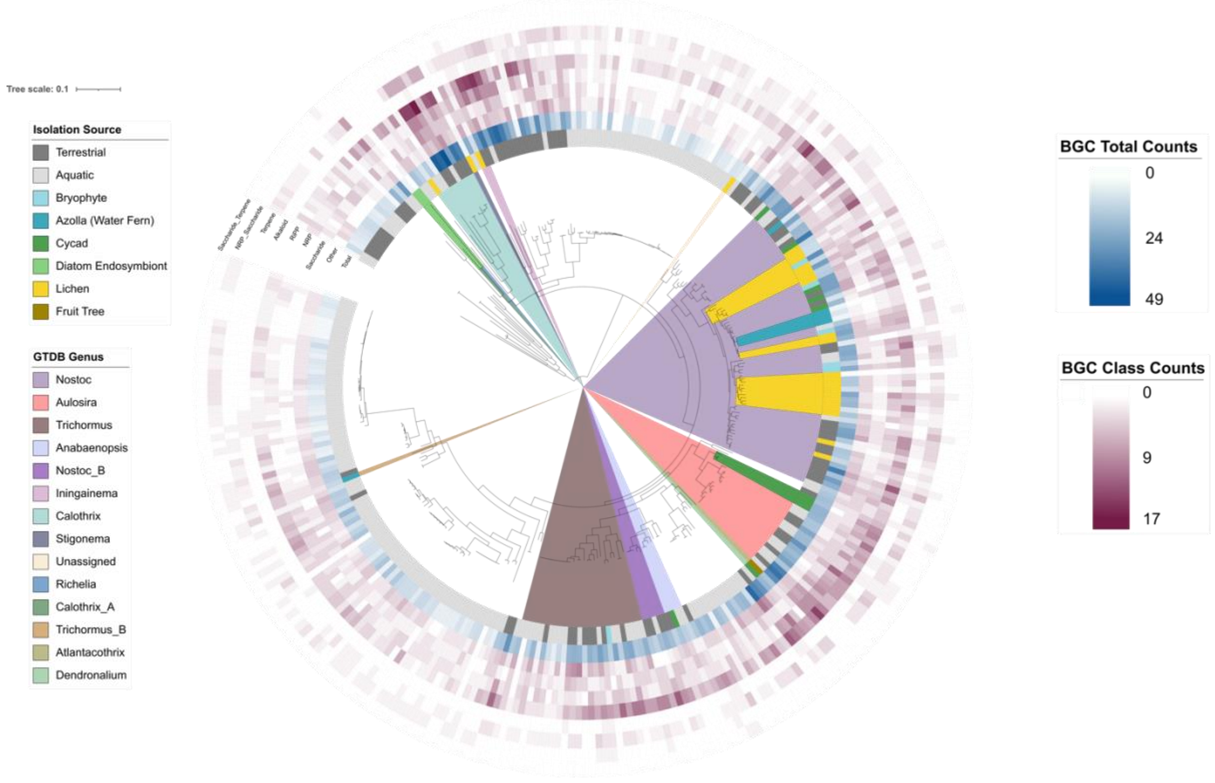


**Figure S6**: Phylogeny generated using concatenated marker genes of genome sequences of strains from family Nostocaceae, rooted with representatives of the order Elainellales, with 1000 bootstraps. Genera with symbiotic lifestyles are highlighted. From inside to outside annotation tracks depict the i) isolation source, and ii) remaining tracks depict the distribution of total counts and counts of individual biosynthetic gene clusters found to be significantly associated with isolation source.

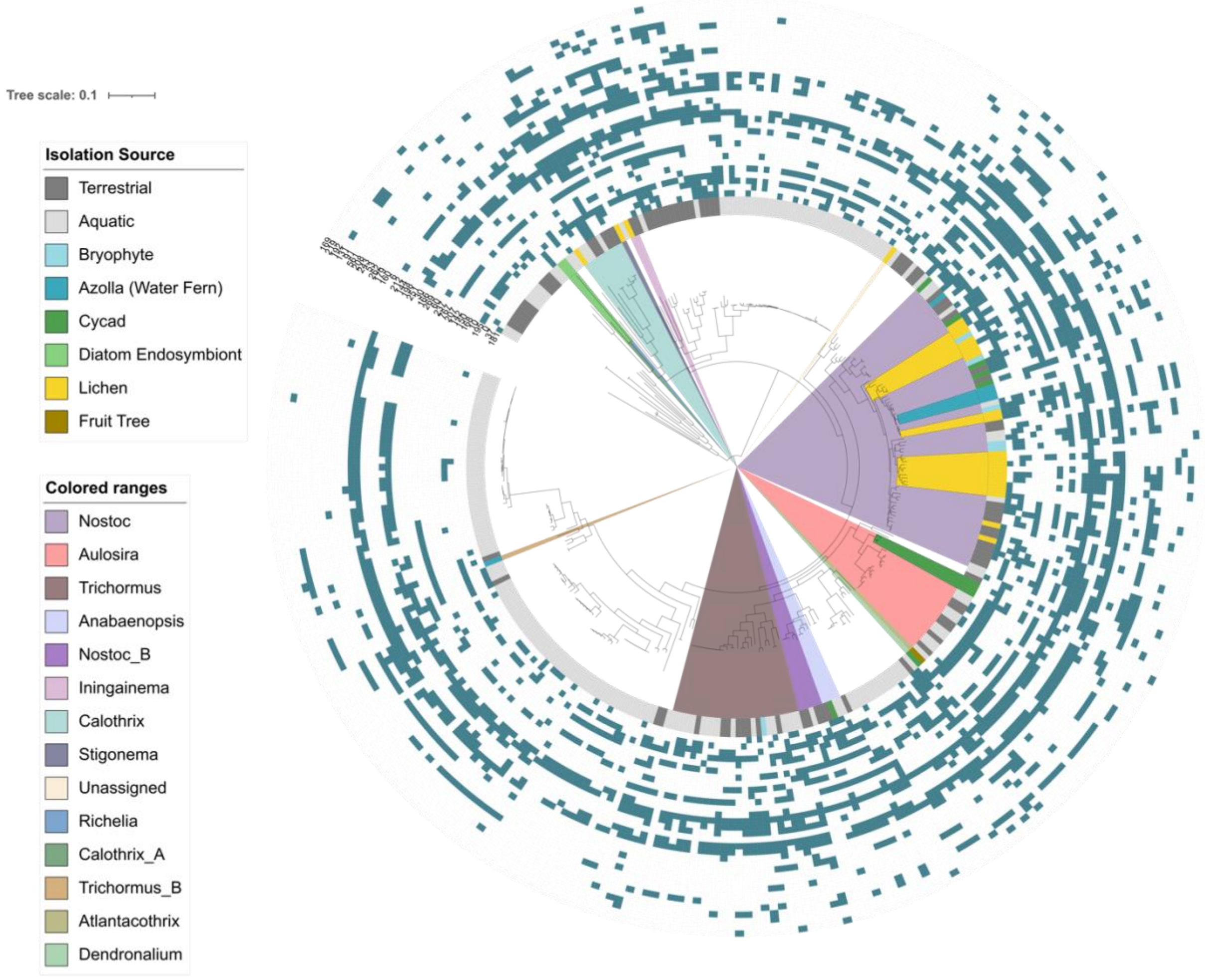


**Figure S7:** Phylogeny generated using concatenated marker genes of genome sequences of strains from family Nostocaceae, rooted with representatives of the order Elainellales, with 1000 bootstraps. Genera with symbiotic lifestyles are highlighted. From inside to outside annotation tracks depict the i) isolation source, and ii) remaining tracks depict the distribution of BGC groups generated through Louvain4 clustering found to be significantly associated with isolation source.
